# A new method for improving extraction efficiency and purity of urine and plasma cell-free DNA

**DOI:** 10.1101/2020.12.31.425003

**Authors:** Selena Y. Lin, Yue Luo, Matthew M. Marshall, Barbara J. Johnson, Sung R. Park, Zhili Wang, Ying-Hsiu Su

## Abstract

This study assessed three commercially available cell-free DNA (cfDNA) extraction kits and the impact of a PEG-based DNA cleanup procedure (DNApure) on cfDNA quality and yield. Six normal donor urine and plasma samples, and specimens from four pregnant (PG) women carrying male fetuses underwent extractions with the JBS cfDNA extraction kit (kit J), MagMAX Cell-Free DNA Extraction kit (kit M), and QIAamp Circulating Nucleic Acid Kit (kit Q). Recovery of a PCR product spike-in, endogenous *TP53,* and Y-chromosome DNA was used to assess kit performance. Nucleosomal-sized DNA profiles varied among the kits, with prominent multi-nucleosomal-sized peaks present in urine and plasma DNA isolated by kits J and M only. Kit J recovered significantly more spike-in DNA compared with kit M or Q (*p*<0.001) from urine, and similar amounts from plasma (*p*=0.12). Applying DNApure to kit M- and Q-isolated DNA significantly improved the amplification efficiency of spike-in DNA from urine (*p*<0.001) and plasma (*p*≤0.013). Furthermore, kit J isolated significantly more Y-chromosome DNA from PG urine compared to kit Q (*p*=0.05). We conclude that DNApure provides an efficient means of improving the yield and purity of cfDNA and minimizing effects of pre-analytical biospecimen variability on liquid biopsy assay performance.

## INTRODUCTION

Liquid biopsies present a minimally invasive or noninvasive approach for detecting circulating cell-free DNA (cfDNA) markers in prenatal genetic testing^1–4^, organ transplant monitoring^5–7^, and precision oncology^8–11^. CfDNA isolated from plasma and urine is often low in quantity and highly fragmented as it is mostly derived from apoptotic cells. CfDNA fragments range predominantly from 130 to 250 bp, reflecting the mono-nucleosomal-sized DNA as the major cfDNA species in plasma^12–14^ and urine^15^, and are even smaller in the case of circulating cell-free fetal DNA (ccffDNA) in maternal urine^4^. Additionally, detecting circulating tumor (ctDNA)^16^ or ccffDNA^4^ presents its own challenges due to the high background of normal/wild-type cfDNA derived from peripheral blood cells or urinary tract cells.

While cfDNA is a promising analyte for liquid biopsy, detection of cfDNA markers is significantly impacted not only by pre-analytical variables such as cfDNA isolation^17^, but also by biospecimen variability. CfDNA isolation platforms, which are primarily column- or magnetic-bead-based, can be impacted by this variability, including the levels of biologically active molecules, overall molecular composition, and the presence of impurities. This phenomenon in turn results in variable DNA yield, quality, and residual impurity content. Urine is known to be a highly complex sample matrix with high inter- and intra-person variability^18^. In an effort to reduce the impact of biospecimen variability on cfDNA isolation and subsequent analytical performance, we developed a PEG-based DNA cleanup step (DNApure) to remove impurities from extracted cfDNA and cell-associated large genomic DNA.

In this study, we compared the analytical performance of two frequently used, commercially available kits, the bead-based MagMAX Cell-free DNA extraction kit and the column-based QIAamp Circulating Nucleic Acid kit, with that of the JBS cfDNA extraction kit, developed by our laboratory, before and after DNApure cleanup. We compared cfDNA fragment size, yield, purity (as assessed by PCR amplification efficiency), and their reproducibility among the three kits in normal donor urine and plasma, and in specimens collected from pregnant (PG) donors carrying a male fetus. Our results demonstrate that the PEG-based DNApure cleanup procedure improves the purity of cfDNA isolated from urine and plasma by all three kits and highlight the importance of obtaining high quality and quantity of cfDNA to improve downstream liquid biopsy applications.

## RESULTS

### Comparison of cfDNA extraction kit performance in urine

Given the importance of obtaining nucleosomal-sized DNA from liquid biopsies, we first compared the overall fragment size distributions in cfDNA isolated from six normal urine donors with the three kits. Because it is known that sex impacts urinary DNA yield^8^, three male and three female donors were included (**Figure 1A**). Representative electropherograms of cfDNA isolates obtained with each kit are compiled for each donor in **Figure 2**. Interestingly, in some donors the DNA size profiles were different among the kits. For example, donors 1 and 6 showed detectable mono- and di-nucleosomal-sized peaks in kit J and M, but not in kit Q isolates. Overall, all three kits were able to isolate mono-nucleosomal-sized DNA, except in male donor #5, whose samples produced low DNA yields from all three kits.

**Figure 1.**
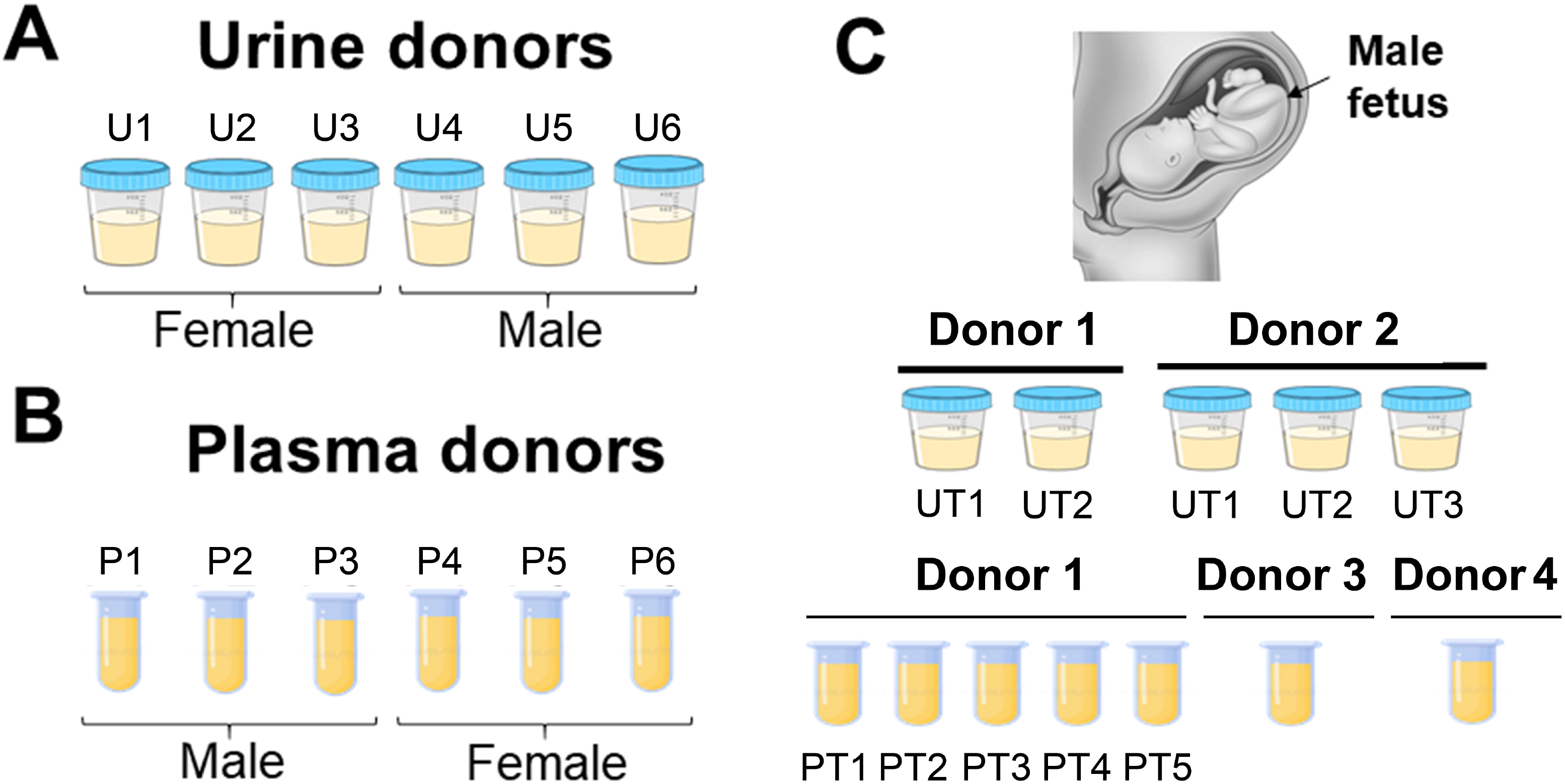
Normal donor specimens used in study. A) Six normal urine donors (U1-6). B) Six normal plasma donors (P1-6). Urine and plasma donors are non-matched. C) Urine and plasma specimens collected from four third-trimester pregnant (PG) donors carrying a male fetus. Donor 1 (UT1, UT2) and 2 (UT1-3) urine was collected at five different timepoints. Donor 1 PG plasma was collected in-house at PT1-PT5 timepoints. Donor 3 and 4 PG plasma was collected by Lee BioSolutions. All urine samples were collected in-house in a final 30-50mM EDTA concentration.

**Figure 2.**
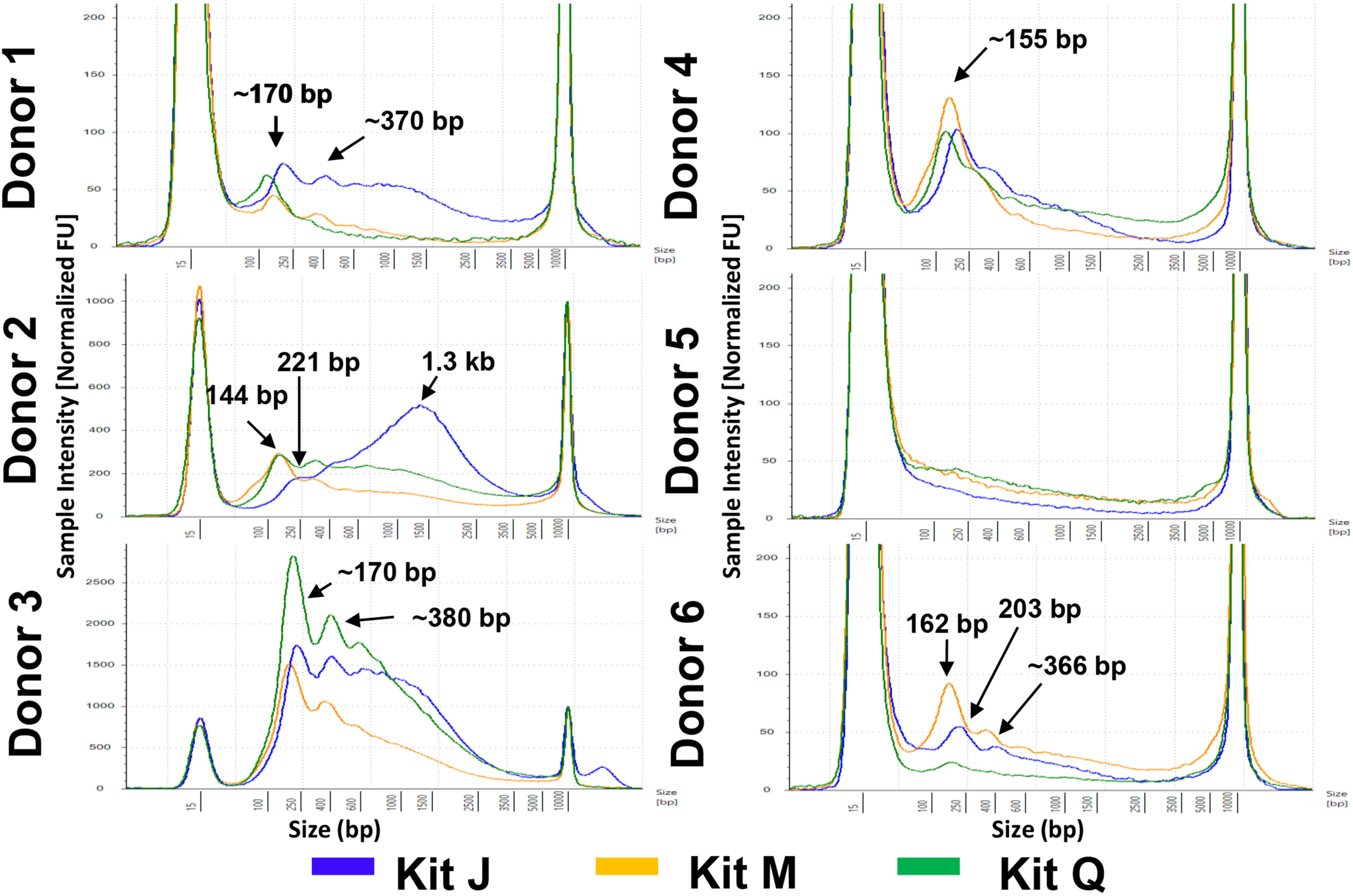
CfDNA fragment size profiles of urine samples extracted from 6 normal donors. Representative profiles are shown for each donor after extraction following the manufacturers’ protocols. DNA input equivalent of 0.08mL urine was loaded onto a D5000 High Sensitivity screentape. TapeStation-software-identified DNA peaks or average (denoted with “~”) peak size are indicated given ± 15% sizing accuracy of screentape. All electropherograms are scaled to sample. Kit J (blue), kit M (orange), and kit Q (green) cfDNA size profiles are indicated by color.

Impurities, salt, or the amount of large genomic DNA in the isolated cfDNA could impact the size distribution and peak heights in the electropherogram analysis. As mono-nucleosomal-sized DNA is the predominant species of cfDNA, we evaluated the efficiency of the three kits in recovering mono-nucleosomal-sized DNA. We utilized a 141 bp double-stranded PCR product spike-in as a mimic of mono-nucleosomal-sized cfDNA to assess recovery efficiency. The PCR product was added into each urine sample at 10^6^ copies/mL. As assessed by qPCR, the spike-in DNA was recovered in a range of 7.8×10^4^ – 3.7×10^6^ copies/mL among the six donors. However, on average, kit J recovered significantly more spike-in DNA compared with kit M or Q (*p*<0.001) (**Supplemental Table 1, Figure 3A**). The apparent spike-in recovery efficiency exceeding 100% in some samples may indicate further spike-in PCR product purification during the extraction process, improving the downstream PCR amplification efficiency.

**Figure 3.**
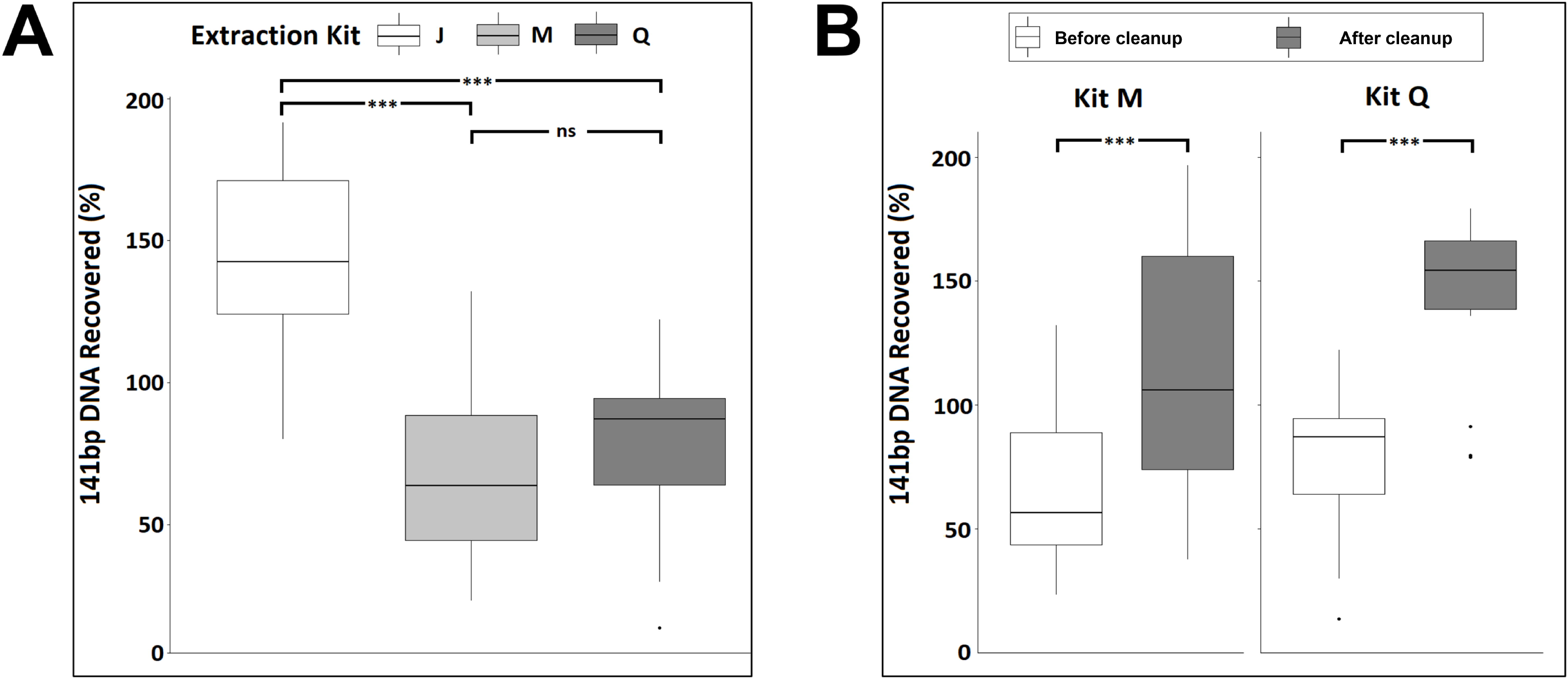
Synthetic 141 bp spike-in DNA recovery from six normal urine donors by three cfDNA extraction kits as measured by qPCR before (A) and after (B) DNApure cleanup. For each kit, triplicate extractions were performed for each of the six urine donors. “Cleanup” indicates samples were further purified using DNApure cleanup. Symbols indicate significance levels of differences between samples under brackets as assessed with Tukey’s post hoc test (ns, >0.1; ***, <0.001).

To better understand if the wide range of spike-in DNA recovery was due to impurities/PCR inhibitors in the eluted urine cfDNA, we measured the protein levels in all extracted urine cfDNA samples and found no detectable protein. Next, we applied the DNApure cleanup to urine DNA isolated with kits M and Q to assess whether it could improve PCR quantitation of the spike-in DNA. As shown in **Figure 3B**, a significant improvement of the PCR amplification efficiency of the spike-in DNA was achieved for both kits (*p*<0.001) after the cleanup. The spike-in DNA recovery rates achieved with the DNApure procedure were comparable among all three kits (*p*=0.482). Next, to assess the yields of urine cfDNA isolated with each kit after DNApure cleanup, we performed qPCR-based *TP53* gene quantification in extracted DNA. As expected, the urine *TP53* levels ranged widely among the donors, from 1.77×10^2^ to 1.67×10^5^ copies/mL (**Supplemental Figure 1**) and did not differ significantly among the kits (*p*=0.23).

### Performance of the JBS cfDNA extraction kit in plasma

To assess if the PEG-based DNApure cleanup can also be applied to plasma cfDNA isolation, we next assessed the performance of the three kits in normal plasma donors (**Figure 1B**), utilizing 2mL of plasma for all extractions, as described in Materials and Methods. Plasma cfDNA isolated from all six donors displayed a prominent mono-nucleosomal-sized peak when isolated with all three kits (**Figure 4).** Four donors displayed prominent di- and tri-nucleosomal DNA peaks in plasma cfDNA samples isolated by kits J and M; however, these peaks were undetectable or reduced in cfDNA isolated from the same donors by kit Q. The recovery of spike-in DNA did not differ significantly (*p*=0.12) among the kits (**Figure 5A**). The PCR amplification efficiency of the recovered spike-in DNA significantly improved after the DNApure cleanup in samples extracted with both kit M (*p*=0.013) and kit Q (*p*<0.001) (**Figure 5B**). Interestingly, protein contamination was only detected in plasma cfDNA samples isolated with kit Q before cleanup (**Supplemental Figure 2)** and was no longer detectable after cleanup (data not shown). Lastly, the *TP53* yield did not differ significantly (*p*=0.61) among the kits (**Supplemental Figure 3**) after DNApure cleanup. Overall, the plasma cfDNA extraction is highly comparable among all three kits after cleanup as PEG-based DNApure cleanup enabled removal of PCR inhibitors and proteins co-purified with plasma cfDNA.

**Figure 4.**
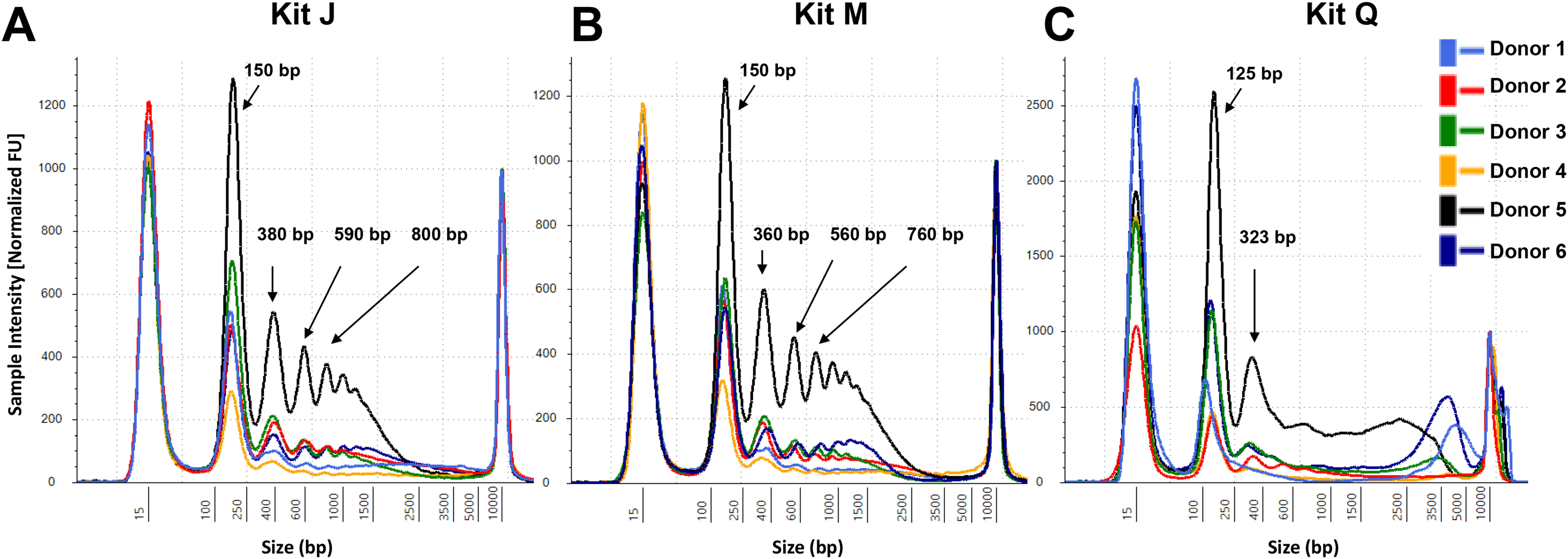
CfDNA size profiles of plasma samples extracted from six normal donors. Representative profiles are shown for each donor. cfDNA extracted with kits M and Q was subjected to the DNApure cleanup procedure prior to TapeStation loading to remove gel impurities. DNA input equivalent of 0.08mL plasma was loaded onto a D5000 High Sensitivity screentape. Averaged TapeStation software-identified DNA peaks are indicated.

**Figure 5.**
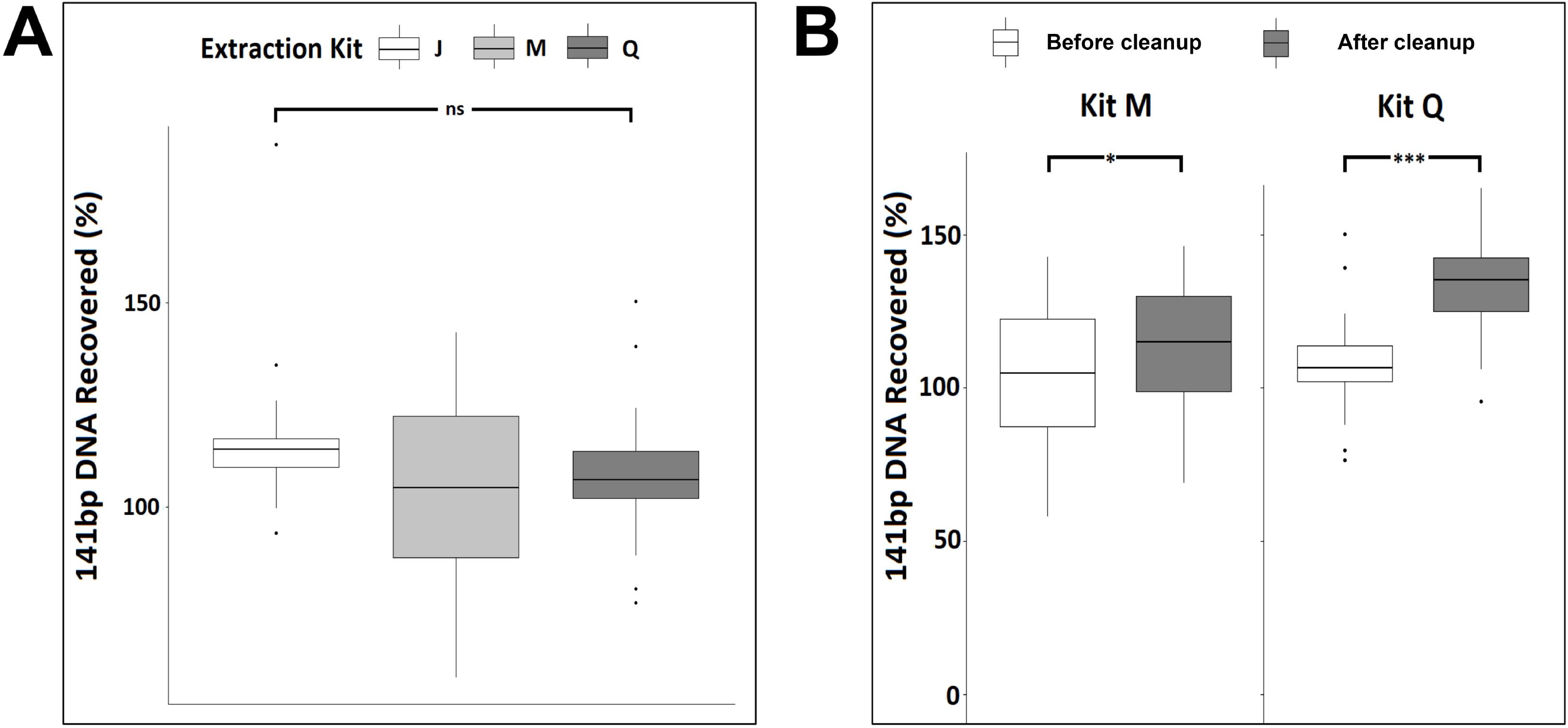
Synthetic 141 bp spike-in DNA recovery from six normal plasma donors by three cfDNA extraction kits as measured by qPCR before (A) and after (B) DNApure cleanup. For each kit, triplicate extractions were performed for each of the six plasma donors. “Cleanup” indicates samples were further purified using DNApure cleanup. Symbols indicate significance levels of differences between samples under brackets as assessed with Tukey’s post hoc test (ns, >0.1; *, <0.05; ***, <0.001).

### Extraction of circulating fetal DNA in liquid biopsies

CcffDNA is released into the maternal circulation and can be detected in urine. Utilizing samples from four PG donors carrying a male fetus (**Figure 1C**), we assessed the efficiency of ccffDNA extraction from urine and plasma samples with three kits according to the manufacturers’ protocols. A total of five urine specimens were collected from two third-trimester PG donors at various timepoints. We found the male Y-chromosome quantities to be significantly higher in kit J isolates compared to kit Q isolates (*p*=0.05), with no significant differences between kits J and M (*p*=1.0) (**Figure 6A**). Due to limited volumes of blood collected, only kits J and Q were evaluated on all seven plasma collections. As shown in **Figure 6B**, Y-chromosome quantities extracted from PG plasma were similar between the two kits.

**Figure 6.**
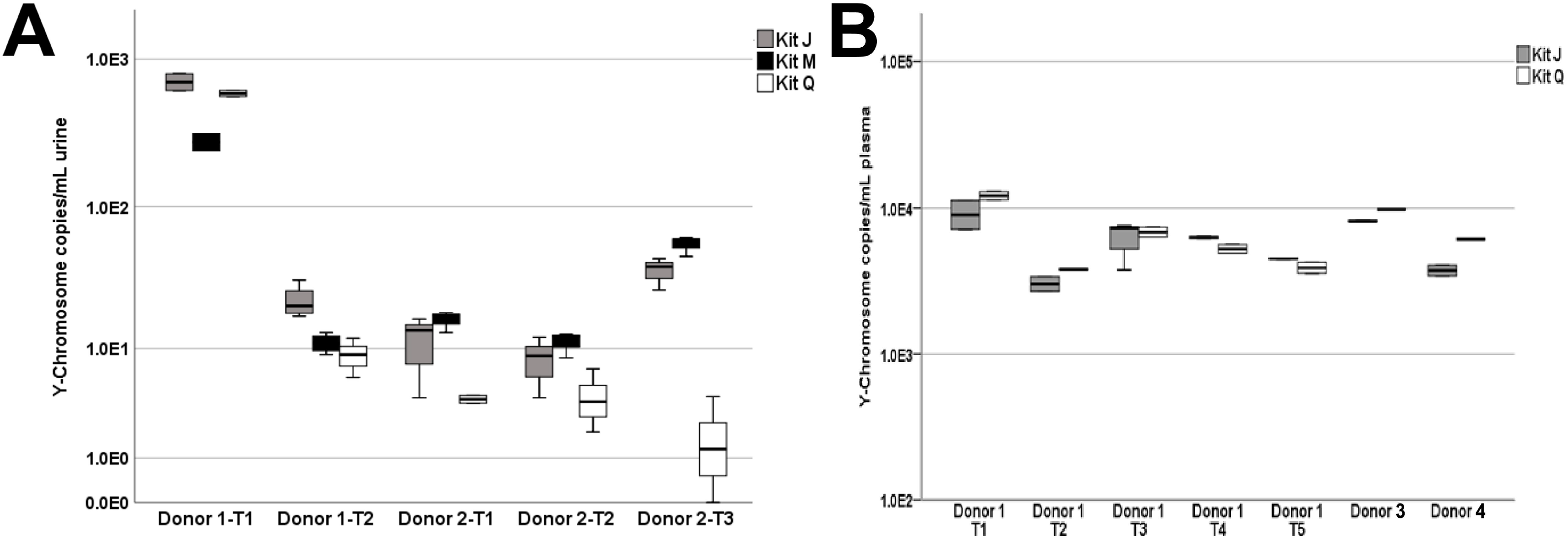
Y-chromosome DNA extracted from liquid biopsy specimens of pregnant donors carrying a male fetus. A) Y-chromosome quantities isolated from urine collected from two donors over five time points (Donor 1: T1 and T2; Donor 2: T1, T2, and T3). B) Y-chromosome quantities isolated from 0.5mL plasma. Donor 1 had matched plasma sampled at times T1 and T2 and subsequently plasma-only at times T3, T4, and T5. Due to limited PG plasma quantities, cfDNA was extracted with kits J and Q only.

## DISCUSSION

In this study, we compared the performance of the urine and plasma cfDNA isolation procedures embodied in three different kits and assessed the impact of the PEG-based DNApure method on cfDNA yields in various specimens. We used Y-chromosome DNA in PG urine or plasma as an unambiguous circulating cfDNA marker to compare the isolation efficiencies of circulation-derived cfDNA. We demonstrated the performance of the DNApure-integrated JBS kit is similar to those of two widely used (one bead-, one column-based) commercial kits for isolating the quantity of cfDNA (measured by less sensitive methods) and successful in extraction of mono- and, frequently, di-nucleosomal-sized DNA. The JBS kit recovered significantly higher amounts of spike-in DNA as measured by qPCR assay, compared to the other kits and extracted higher amounts of ccffDNA. The PEG-based DNApure cleanup can effectively remove impurities present in isolates obtained with the other kits, thus improving their downstream PCR amplification. This finding is highly significant since PCR amplification is a key methodology used in almost all liquid biopsy assay platforms.

The comparison of the three kits across six normal urine donors and their replicates demonstrates the need for an extraction method capable of robustly handling a wide range of biospecimen variability. Biospecimen variability can stem from factors such as sex, disease type, and collection method and time. In this study, three male and three female urine donors were included as female urine is known generally to contain higher amounts of cfDNA than male urine due to increased release of cfDNA from the urinary tract. Consistent with this notion, samples from male donor #5 produced the lowest yields of urine cfDNA among all three kits. Interestingly, PG donor 1 T1 and T2 urine samples were collected less than one week apart, but contained vastly different amounts of Y-chromosome ccffDNA, suggesting multi-day specimen collection may be needed to increase detection sensitivity of the cfDNA of interest. Biospecimen variability among donors is further highlighted in the cfDNA size profiles of plasma samples extracted with the three kits. The visualization of nucleosomal-sized DNA by TapeStation is known to be impacted by salt/contaminants as well as the presence of large genomic DNA fragments. It is therefore possible the reduced di-and tri-nucleosomal-sized DNA peaks we observed in half of the plasma cfDNA samples extracted by kit Q may be caused by impurities. It is not surprising that PEG-based cleanup was able to remove proteins and PCR inhibitors in cfDNA extracted from liquid biopsy samples (Figures 3, 5, and Supplemental Figure 2), as PEG-based solutions have been used in previously developed DNA extraction and purification procedures^19,20^. The similar total amounts of cfDNA after cleanup, as assessed by *TP53* qPCR, obtained with all three kits are consistent with this explanation. Overall, these observations highlight the value of the PEG-based DNApure cleanup in minimizing the impact of biospecimen variability on DNA isolation and, consequently, procuring high-quality cfDNA from liquid biopsy specimens.

As cfDNA in urine is less concentrated than in plasma^21^, and large volumes of urine are often readily available, we developed our urine DNA isolation method to accommodate large input volumes (up to 50mL), whereas a majority of commercial urine cfDNA extraction kits cannot process volumes more than 4 mL at a time. Similar to the other two kits, JBS urine and plasma cfDNA extraction can be automated using the JBS JPurX-S200 instrument to reduce variation due to manual processing. While only normal donors were included in this study, extraction of urine cfDNA from donors with various diseases, such as cancer^22,23^ and infection-related conditions^22^, may be hampered by the even greater biospecimen variability associated with such pathologies. The effects of this variability on the performance of any new cfDNA isolation method will need to be thoroughly assessed.

In summary, this study brings forth the importance of cfDNA isolation methodology in achieving high cfDNA quality and the potential value of the PEG-based DNApure procedure in mitigating the impact of biospecimen variability on analytical performance. By improving the quality and yield of isolated cfDNA, downstream analyses can also be significantly improved as well as standardized, enhancing liquid biopsy utility and facilitating the adoption of cfDNA-based assays in the clinic.

## MATERIALS AND METHODS

### Urine and plasma sample collection

All human specimens used in this study were either purchased from BioIVT^®^ (Westbury, NY, USA), Lee Biosolution (Maryland Heights, MO, USA), or archived, de-identified, and collected from other studies, as described previously^15^, thus the only patient information obtained was sex (**Supplemental Table 2**). Frozen human urine (50mL) was collected from healthy female (n=3) and male (n=3) volunteers and mixed with EDTA to achieve 30-50mM final EDTA concentration. Normal donor blood collected in K_2_EDTA tubes was obtained from BioIVT^®^. Plasma was separated from whole blood by centrifugation at 2,800 x g for 20 min and filtered (0.2μm pore size) in 2.0mL aliquots. Plasma and urine specimens were collected from four PG females in their third trimester carrying a male fetus. The first PG donor had matched urine and plasma collected at two different times, T1 and T2, and three subsequent plasma-only collections (PT3, PT4, and PT5). PG donor 2 provided only urine samples collected at three different times, T1, T2, and T3. PG donors 3 and 4 had only plasma obtained by Lee Biosolution. In this case, blood was collected by venipuncture, centrifuged at 3,500 RPM, then the plasma was pipette off and stored at −20°C. Specimens used in this study are shown in **Figure 1**. All experiments were performed in accordance with relevant guidelines and regulations.

### CfDNA extractions

CfDNA was extracted from normal urine and plasma samples in 3-4 replicates using each of the three cfDNA extraction kits following the manufacturers’ protocols: the JBS DNA extraction kit, which includes PEG-based DNApure cleanup (kit J; cat# JBS-08872 for urine or cat# JBS-08874 for plasma; JBS Science, Doylestown, PA, USA), the MagMAX Cell-Free DNA Extraction (kit M; cat# A29319, Thermo Fisher Scientific, Waltham, MA, USA), and the QIAamp Circulating Nucleic Acid kit (kit Q; cat# 55114, Qiagen, Germantown, MD, USA).

Each urine/plasma specimen was pooled, mixed well, and aliquoted for extraction. Extraction was done according to the manufacturers’ protocols utilizing 3.0mL urine or 2.0mL plasma aliquots in triplicate. Kit M and Q urine and plasma samples were ultra-centrifuged at 16,000 x g for 10 minutes at 4°C prior to cfDNA extraction. Aliquots were immediately extracted or frozen at −20°C until cfDNA extraction was performed. Due to the low amounts of fetal DNA found in maternal urine, PG urine samples were extracted using up to 4.0mL urine input. Due to the limited quantities of PG plasma obtained from Lee Biosolution, extraction of 0.5mL plasma aliquots was performed for kit J and Q assessment only.

### Synthetic 141 bp spike-in DNA

Normal urine and plasma biopsies were spiked at 10^6^ copies/mL with a PCR-synthetic 141 bp double-stranded DNA fragment prior to cfDNA extraction. The synthetic spike-in DNA was quantified using the JBS Artificial Spike-in DNA Quantification kit according to the manufacturer’s specification and the recovered copies were estimated in each cfDNA sample. We calculated spike-in recovery as:

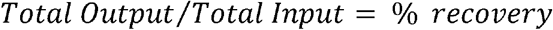

### Size assessment

We visualized cfDNA profiles on High Sensitivity D5000 ScreenTapes on the TapeStation 4200 system (Agilent Technologies, Santa Clara, CA, USA). The equivalent of 0.08mL urine or plasma was visualized for each specimen.

### Quantification assays

*TP53* gene quantity was measured by qPCR as previously described^24^ and Y-chromosome DNA was measured using the Y-Chromosome DNA quantification kit (JBS Science). To assess the amounts of residual protein impurities after cfDNA isolation, protein concentration of each sample was determined using the Qubit Protein Assay kit (cat# Q33211, Thermo Fisher Scientific) with an input equivalent of 0.08mL urine and 0.08-0.20mL plasma. Quantitative PCR assays were performed in duplicate on the LightCycler 480 platform (Roche, Indianapolis, IN, USA).

### Statistical Analyses

We tested data for normality using the Shapiro-Wilk test and for homogeneity of variance using Levene’s test. We tested the hypothesis of no difference among DNA extraction kits J, M, and Q in recovery of synthetic 141 bp DNA and in recovery of *TP53* DNA using analysis of variance (ANOVA) with extraction kit as a fixed effect, or the Kruskal-Wallis test among extraction kits when parametric assumptions were not met. For post hoc comparisons among DNA extraction kits, we used Tukey’s post hoc comparison for ANOVA procedures. We tested the hypothesis of no difference in recovery of synthetic 141 bp DNA in samples extracted with kits M and Q before and after conducting the DNApure cleanup procedure using the paired t-test or the Wilcoxon signed-rank test when parametric assumptions were not met. For all statistical tests we used α= 0.05 to determine whether differences were significant. All analyses were conducted in R v3.6.1 (R Development Core Team, 2013).

## Data Availability Statement

The data generated and analyzed during this study are available from the corresponding author on reasonable request.

## Acknowledgements

We thank Dr. Dmitry Goryunov and Jackie Kuo for their support in proofreading and editing of this manuscript.

## Author contributions

S.Y.L and Y.S. designed the study. S.Y.L, Y.L., M.M.M., Z.W., and B.J.J performed the experiments. S.YL., Y.L., M.M.M., B.J.J., Z.W., and S.R.P. performed the data analysis. S.Y.L. and Y.S wrote the manuscript. All authors reviewed the manuscript.

