## Supplemental Data for "A new method for improving extraction efficiency and purity of urine and plasma cell-free DNA"

**Figure S1.** *TP53* DNA concentrations in cfDNA isolated from six normal urine donors per kit.

**Figure S2.** Protein contamination in cfDNA extracted from plasma (2mL) of six normal donors by three cfDNA extraction kits as measured by Qubit total protein assay.

**Figure S3.** *TP53* DNA concentrations isolated from six normal plasma donors per kit.

**Table S1.** Statistical results of Tukey’s post hoc comparisons for recovery of synthetic 141 bp DNA from 3mL urine samples with kits J, M, and Q.

**Table S2.** Liquid biopsy donor sample information**.**

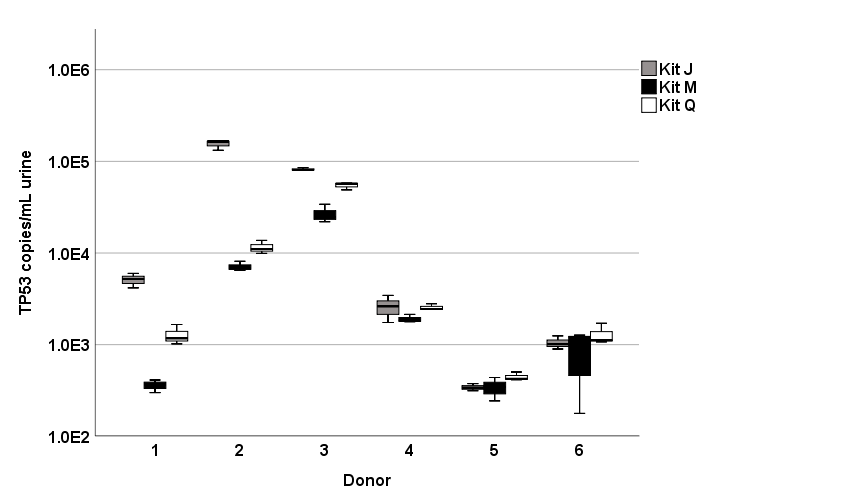

**Figure S1. *TP53* DNA concentrations in cfDNA isolated from six normal urine donors per kit.** Average *TP53* concentrations of three extraction replicates for each donor are shown.

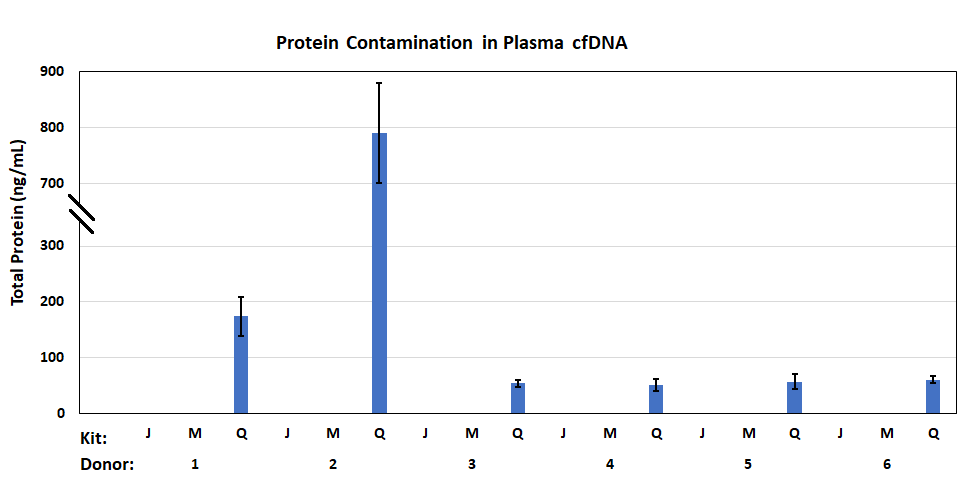

**Figure S2. Protein contamination in cfDNA extracted from plasma (2mL) of six normal donors by three cfDNA extraction kits as measured by Qubit total protein assay.** DNA input equivalent of 0.2mL of plasma was used.

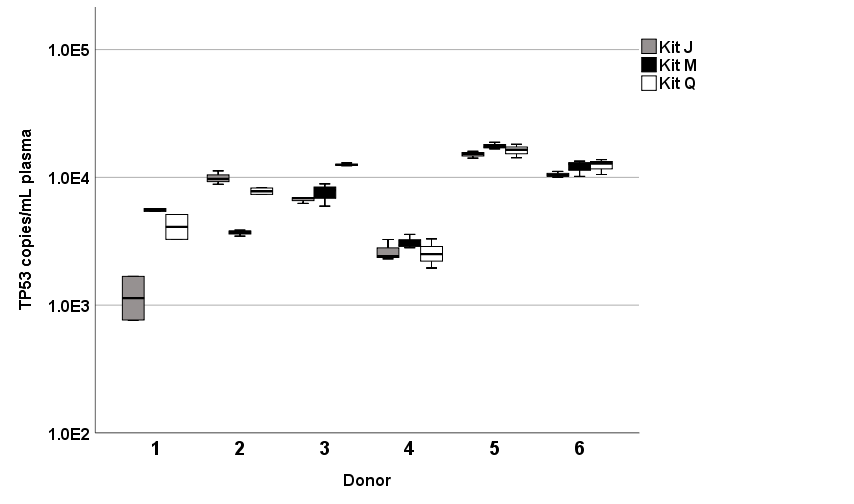

**Figure S3. *TP53* DNA concentrations isolated from six normal plasma donors per kit.** Average *TP53* copies/mL concentrations are shown per donor from three extraction replicates per kit.

**Table S1. Statistical results of Tukey’s post hoc comparisons for recovery of synthetic 141 bp DNA from 3mL urine samples with kits J, M, and Q.**

| Contrast | Estimate | 95% CI | d.f. | t | p-value |
| --- | --- | --- | --- | --- | --- |
| J - M | 0.75 | 0.63-0.86 | 44 | 11.72 | <0.001 |
| J - Q | 0.65 | 0.51-0.79 | 44 | 9.94 | <0.001 |
| M - Q | -0.1 | -0.21-0.02 | 44 | -1.61 | 0.25 |

95% CI, 95% confidence interval; df, degrees of freedom; t, t-value.

**Table S2. Liquid biopsy donor sample information.**

| Biopsy | Donor # | Sex | Biopsy extraction volume (mL) |
| --- | --- | --- | --- |
| Urine | 1 | F | 3 |
| Urine | 2 | F |  |
| Urine | 3 | F |  |
| Urine | 4 | M |  |
| Urine | 5 | M |  |
| Urine | 6 | M |  |
| Plasma | 1 | M | 2 |
| Plasma | 2 | M |  |
| Plasma | 3 | M |  |
| Plasma | 4 | F |  |
| Plasma | 5 | F |  |
| Plasma | 6 | F |  |
| PG Urine | Donor 1 T1 | F | 4mL |
| PG Urine | Donor 1 T2 | F | 4mL |
| PG Urine | Donor 2 T1 | F | 4mL |
| PG Urine | Donor 2 T2 | F | 4mL |
| PG Urine | Donor 2 T3 | F | 4mL |
| PG Plasma | Donor 1 T1 | F | 0.5mL |
| PG Plasma | Donor 1 T2 | F | 0.5mL |
| PG Plasma | Donor 1 T3 | F | 0.5mL |
| PG Plasma | Donor 1 T4 | F | 0.5mL |
| PG Plasma | Donor 1 T5 | F | 0.5mL |
| PG Plasma | Donor 3 | F | 0.5mL |
| PG Plasma | Donor 4 | F | 0.5mL |
